# How Does Perceptual Discriminability Relate to Neuronal Receptive Fields?

**DOI:** 10.1101/2022.12.21.521510

**Authors:** Jingyang Zhou, Chanwoo Chun

## Abstract

Perception is an outcome of neuronal computations. Our perception changes only when the underlying neuronal responses change. Because visual neurons preferentially respond to adjustments in some pixel values of an image more than others, our perception has greater sensitivity in detecting change to some pixel combinations more than others. Here, we examined how perceptual discriminability varies to arbitrary image perturbations assuming different models of neuronal responses. In particular, we investigated that under the assumption of different neuronal computations, how perceptual discriminability relates to neuronal receptive fields – the change in pixel combinations that invokes the largest increase in neuronal responses. We assumed that perceptual discriminability reflects the magnitude of change (the L2 norm) in neuronal responses, and the L2 norm assumption gained empirical support. We examined how perceptual discriminability relates to deterministic and stochastic neuronal computations. In the case of deterministic neuronal computations, perceptual discriminability is completely determined by neuronal receptive fields. For multiple layers of canonical linear-nonlinear (LN) computations in particular (which is a feed-forward neural network), neuronal receptive fields are linear transforms of the first-layer neurons’ image filters. When one image is presented to the neural network, the first-layer neurons’ filters and the linear transform completely determine neuronal receptive fields across all layers, and perceptual discriminability to arbitrary distortions to the image. We expanded our analysis to examine stochastic neuronal computations, in which case perceptual discriminability can be summarized as the magnitude of change in stochastic neuronal responses, with the L2 norm being replaced by a Fisher-information computation. Using a practical lower bound on Fisher information, we showed that for stochastic neuronal computations, perceptual discriminability is completely determined by neuronal receptive fields, together with how responses co-variate across neurons.

## 1 Introduction

To quantify a visual neuron’s response, we may start by characterizing its receptive field. Traditionally, receptive field has been regarded as a neuron’s most preferred image pattern (i.e. the image pattern that triggers the highest neuronal firing rate) (Hartline [1938], Kuffler [1953], Hubel and Wiesel [1959, 1962], Spinelli [1966]). To search for a visual neuron’s most preferred image can be challenging in general, especially when images consist of numerous pixels, and the neuron non-linearly responds to pixel combinations in complicated ways (e.g. when the neuron is deep in the visual cortex). In practice, receptive fields are commonly measured as the pattern super-imposed onto a reference image that maximally increases a neuron’s firing response (e.g. Goldin et al. [2022]). In this type of receptive field measurement, a neuron is presented with a reference image, and many patterns of white noise superimposed onto the reference (e.g. Sakai et al. [1988], Chichilnisky [2001]). Averaging across the noise patterns that trigger extra spikes from the neuron (as compared to when the reference is presented alone), we obtain a pattern, or a distortion to the reference image that is effective in increasing neuronal spiking responses. This alternative definition of receptive field is *local*, because a neuron’s receptive field under this definition can be different when measured using different reference images. Empirical evidence supports that neurons in retina and in visual cortices exhibit reference-dependent receptive fields (Barlow et al. [1957], Spinelli [1966], Wielaard and Sajda [2005], Bertalmio et al. [2020], Goldin et al. [2022]).

The study of perception, on the other hand, begins by examining perceptual changes (Fechner [1860]). Typically, perceptual changes are also measured using an image pattern super-imposed onto a reference image. Experimentally, this image pattern is scaled up and down in amplitude until it becomes just perceivable (Green and Swets [1966], Kingdom and Prins [2009]). The just-perceivable scale for a super-imposed image pattern is called the pattern’s perceptual threshold. The inverse of threshold, perceptual discriminability, measures how much perceptual change is induced by scaling the pattern by an (image) unit. Historically, perceptual studies focused on quantifying thresholds using pre-specified image patterns (e.g. adding brightness or contrast to an image). With tools recently developed from computational neuroscience and machine learning, perceptual scientists started to inquire comparatively, which patterns super-imposed onto a reference can make larger difference to perception, e.g. Ahumada et al. [2006], Berardino et al. [2018], da Fonseca and Samengo [2018], Martinez-Garcia et al. [2018].

When pixel values of an image smoothly vary, typically neuronal responses also smoothly vary, and perceptual discrimination was hypothesized to quantify the extent of change in neuronal responses (von Helmholtz [1883], Schrödinger [1920], Vos [1979], Knoblauch and Maloney [1995], Zhou et al. [2021]). Early visual neuroscience literature had a keen interest to relate perceptual discrimination to neuronal response changes. In particular, the literature focused on distorting a reference image along one direction, and comparing discrimination performance of a human or a macaque, to discrimination of individual neurons along that direction (e.g. Spinelli [1966], Britten et al. [1992], Zohary et al. [1990], Liu and Newsome [2005], Geisler et al. [1992]). In this paper, we approached to understand the relationship between neuronal responses and perceptual discrimination from a different angle. We focused on analyzing how to relate perceptual discrimination to neuronal responses using both high-dimensional reference images, and high-dimensional image distortions. We do so because neuronal receptive fields have been quantified using high-dimensional image distortions, but perceptual discriminability has been commonly studied using a single up to a few dimensions, e.g. Macmillian and Creel [1991]. We believe that this mismatch in dimensionality created an artificial barrier to connecting perception to neuronal responses, and has been preventing the field of perception from utilizing high-dimensional statistical tools to expand the range of questions that one can ask. One important goal of visual neuroscience is to understand how neuronal responses contribute to perception, and in this paper, we take a small but important step, by walking through how to understand the connection between the local linear approximation of neuronal responses (receptive field) and local linear approximation of perception (perceptual discrimination). Our setting and analyses are general, and apply to arbitrary number of image pixels and visual neurons.

## 2 Results

To begin our analysis, we make a simplifying assumption that neuronal responses are linear and deterministic, and we examine the perceptual consequence of these simplified neuronal responses (section 2.1). We gradually build towards examining the perceptual consequence of more realistic neuronal responses. In section 2.2, we analyze how linear-nonlinear (LN) neurons’ responses contribute to perception; in section 2.3, we study the perceptual effect of cascaded LN computations. In the last Results section (2.4), we take one step further by making neuronal responses stochastic, and we analyze how perceptual discrimination stems from stochastic neuronal responses.

### 2.1 Linear neuronal responses

In this section, we first link a single linear neuron’s receptive field to perceptual discriminability, before examining the perceptual effect of population responses.

#### How does perceptual discriminability relate to a single linear neuron’s receptive field?

We list some definitions to begin our analysis, and all vectors throughout the paper, unless specified otherwise, are column vectors. We use vector s = [*s*_1_, *s*_2_,…,*s_p_*]^⊤^ to denote an image s with *p* number of pixels, and *s_i_* indicates the *i^th^* pixel’s brightness. Empirically, many visually activated neurons prefer some set of pixel combinations more than others. For example, some cells in primary visual cortex vigorously respond to grating patterns confined to a spatial location (e.g. Hubel and Wiesel [1959], Ringach [2004]), and ganglion cells in retina are known to prefer a spot of light increment surrounded by light decrement, and vice versa (e.g. Barlow [1953], Kuffler [1953]). We use f = [*f*_1_, *f*_2_,…,*f_p_*]^⊤^ to denote a neuron’s image filter, and we assume that the filter has unit length (||f|| = 1) for simplicity of analysis. For the rest of this section, we further assume that the neuron linearly responds to images. For an image s, the linear neuron’s response can be expressed as:

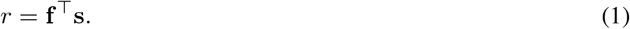

Response *r* increases either when s increases in amplitude (‖s‖ becomes larger or s as an image becomes brighter), or when the pattern of s becomes more similar to the neuron’s filter f. Among all images with a fixed amplitude (e.g. ‖s‖ = 1), the linear computation is reduced to a comparison of angular similarity between filter f and s: f^⊤^s = cos(*θ*), and *θ* is the angle between f and s. Neuronal response is the largest (*r* = 1) when the two vectors are aligned, and the two images share the same pattern. Neuronal response is 0 when the two vectors are orthogonal to each other (see Figure 1).

**Figure 1:**
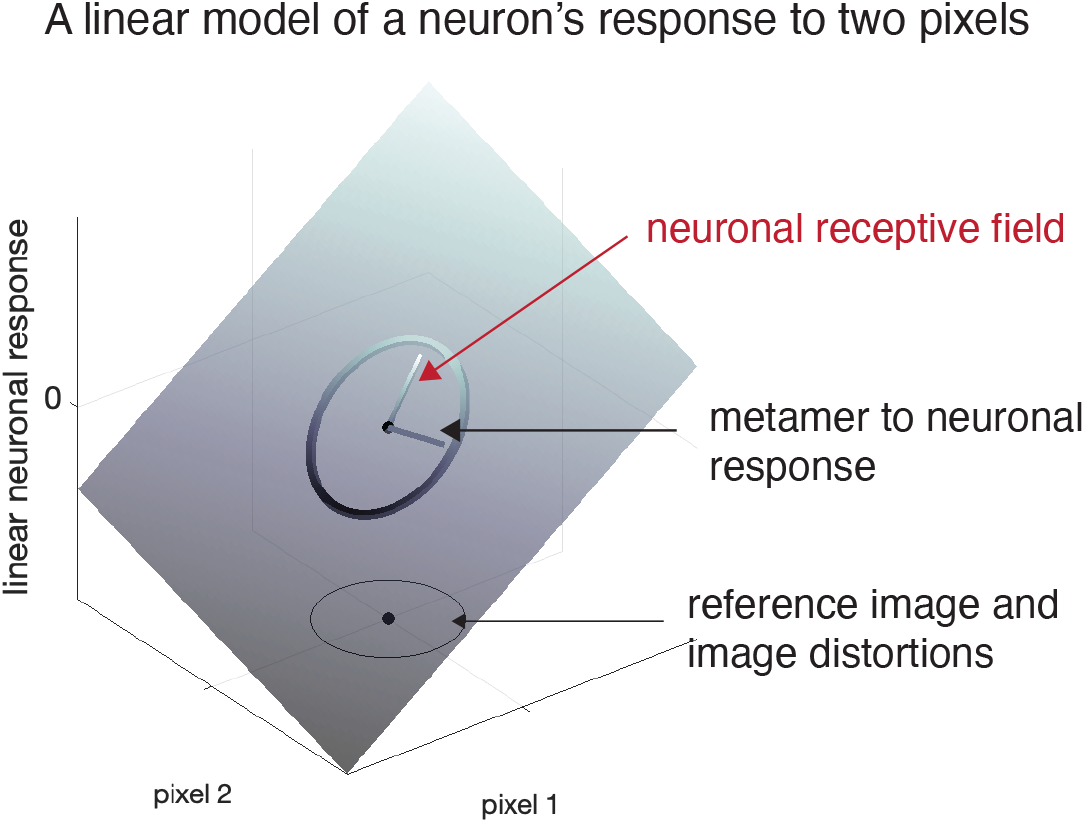
A single neuron’s linear response to two-pixel images. Here, we plotted a neuron’s linear response surface, and its response to a reference image (black dot in the image space), and to distortions of the reference image (the circle in the image space). Across all equi-length image distortions, one distortion direction induces the largest neuronal response increment, and this direction is defined as the neuron’s receptive field. We also plotted an image distortion direction that induces no neuronal response change, and this image distortion is defined as a metamer to the neuron’s response. For a single neuron, perceptual discriminability corresponds to the magnitude of its response change (i.e. the extent of change in the z-axis).

Now we consider a unit-length image pattern **∊** added on top of s, and ‖***∊***‖ = 1. If (s + ***∊***), compared to other patterns ***∊***\* added to s, triggers the largest neuronal response increment, we define ***∊*** as the neuron’s *receptive field* (normalized to unit-length) at reference s. How do we find a linear neuron’s receptive field? To do so, we first need a quantification of stimulus-induced response change. If s only consists of one pixel (s = *s*), to quantify stimulus-triggered response change, we take a derivative of response *r* with respect to *s*. When reference s consists of more than one pixel, we take a similar approach, by computing the directional derivative of *r* with respect to stimulus change along direction ***∊***. The resulting directional derivative is a scalar that quantifies the response change, like in the single-pixel case (see Supplement 4.1). The directional derivative *d_∊_*(s) of a linear neuron has the form:

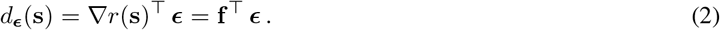

For general neuronal computations, directional derivative compares the response Jacobian ∇*r*(s)^⊤^ to the image distortion **ε**. Neuronal response increment is the largest when ***∊*** and ∇*r*(s) are aligned, and ***∊*** = ∇*r*(s)/‖∇*r*(s)‖. For general neuronal computations, this ***∊*** is the neuron’s receptive field (normalized) at reference image s. For linear computations, in particular, because ∇*r*(s) = f and f is a unit vector, the directional derivative compares the angular similarity between the neuron’s filter f and the image distortion ***∊***. Response increment is the largest when image distortion is aligned to the neuron’s filter, and f = ***∊***. A linear neuron’s receptive field stays the same across all reference images. This is because the response Jacobian (∇*r*(s)^⊤^ = f^⊤^) does not depend on the reference. For general nonlinear computations, a neuron’s receptive field changes with reference images, and is generally distinct from the neuron’s image filter. We will see many examples illustrating this in later sections.

To relate a single neuron’s receptive field to perception, we first make a simplifying assumption that perception is the consequence of one neuron’s computation. We relate the neuron’s computation to the best-developed perceptual measure – perceptual threshold, and its inverse (1/threshold), perceptual discriminability. Experimentally, threshold quantifies how much we need to scale the amplitude of an image distortion in order to induce a unit of perceptual change. Discriminability measures the inverse, how much perceptual change can be triggered by a unit of image distortion (Green and Swets [1966], Kingdom and Prins [2009]). For the rest of this paper, we relate neuronal computations to perceptual discriminability, because it is more natural – like previously we took directional derivatives to quantify image-dependent neuronal response change, discriminability quantifies image-dependent perceptual change.

Discriminability is a non-negative number that summarizes the extent of perceptual change. When a reference image s is perturbed along ***∊***, *d*_∊_(s) denotes the vector of change in neuronal response. We define *perceptual discriminability P_∊_*(s) as the square-root of the magnitude of *d_∊_*(s). In the case of a single neuron, adding or subtracting its receptive field from a reference image is the most perceivable image distortion to the reference. We use the following equations to summarize the relationship between a single neuron’s receptive field ∇*r*(s), perceptual discriminability *p_∊_*(s) and image distortions ***∊***:

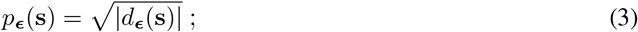

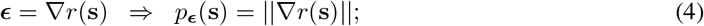

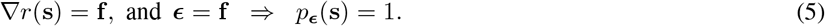

Equation 4 holds for general neuronal computations. For linear neuronal computations (Equation 5), when ∇*r*(s) = f, and when image distortion ***∊*** is aligned to the neuron’s filter, discriminability *p_∊_*(s) is 1, which is the largest discriminability achievable by a linear neuron. Alternatively, if we add or subtract an image pattern ***∊*** that is perpendicular to the neuron’s receptive field, ∇*r*(s)^⊤^ ***∊*** = 0, and the image distortion induces no change in neuronal response (Figure 1). These non-effective image distortions are called *metamers*, e.g. Wandell [1995], Freeman and Simoncelli [2011]. If a neuron’s receptive field is localized, any image distortion out of the spatial extent of the neuron’s activation region is a metamer to the neuron’s response. Metamers can also be patterns within a neuron’s activation region, e.g. Freeman et al. [2015]. An image distortion that is a metamer to neuronal responses should also be a metamer to perception.

#### How does perceptual discriminability relate to a population of linear neurons’ receptive field?

In general, perception results from the computation of a population of neurons (e.g. Paradiso [1988], Seung and Sompolinsky [1993], Kanitscheider et al. [2015], Dalgleish et al. [2020]). Suppose a population consists of *n* neurons, with *k^th^* neuron’s image filter denoted as f_*k*_, and ‖f_*k*_‖ = 1. The population’s filters can be summarized using a *p* × *n* matrix *F*, and *F* = [f_1_, f_2_,…,f_*n*_]. Each neuron in the population linearly converts an image to a single response number *r_k_*, and a column vector *r* summarizes the linear population’s response:

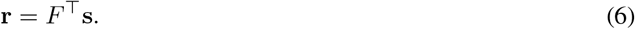

When a reference image s is distorted along **∊**, the response of the population changes, and the change can be summarized using a directional derivative d_*∊*_(s), which is an *n*-dimensional vector in this case:

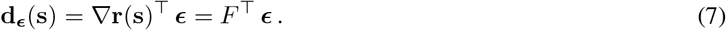

∇*r*(s)^⊤^ is the Jacobian matrix that captures each neuron’s response change. We define ∇*r*(s) as the neuronal population’s *receptive field*. A linear population’s filter *F* coincides with its receptive field.

Distorting image s along different **∊** directions, the populational response may adjust in amplitude, or direction, or both. We highlight two special neuronal populations here to guide our population analysis. First, if all neurons in the population share a single filter (f_*k*_ = f), any image distortion ***∊*** may change the amplitude of the populational response vector, but not its direction (see Supplement 4.2). Alternatively, if the population consists of *p* neurons, all of which have orthogonal filters (so *F* is an orthonormal matrix), different image distortions may only change the direction of the populational response, but not its amplitude (see Supplement 4.2). In general, neuronal filters are probably neither completely parallel nor orthogonal, so for arbitrary image distortions, populational response generally changes both in amplitude and in direction.

To examine the perceptual implication of change in population response, we need to relate d_*∊*_(s) to perceptual discriminability, which is a single non-negative scalar. There are many ways to convert a vector d_*∊*_(s) into a nonnegative number, and in Discussion, we will examine the perceptual consequence of this conversion via different *L^p^* norms: ‖d_*∊*_(s)‖^*p*^. Here, and for the rest of the Results, we assume discriminability as the *L*^2^ norm of the population response change, *p_∊_*(s) = ‖d_*∊*_(s)‖. This is because *L*^2^ norm gained empirical support (Poirson and Wandell [1990], Knoblauch and Maloney [1995]). Additionally, *L*^2^ norm has a simple geometrical interpretation: under the *L*^2^ norm assumption, discriminability measures when a stimulus is perturbed, how much distance is traversed on the manifold of neuronal responses.

In general, perceptual discriminability predicted from a neuronal population has the following expression:

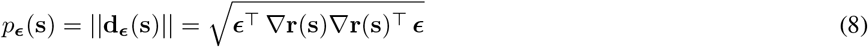

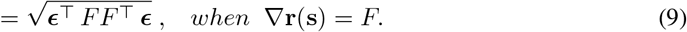

For a linear population, ∇*r*(s) = *F*, and perceptual discriminability can be directly predicted from the population filters. The set of equations above show that perceptual discriminability is *completely* determined by the population receptive field ∇*r*(s). At one reference image s, perceptual discriminability to any image perturbation is determined by the discrimination matrix *J*(s) = ∇*r*(s)∇*r*(s)^⊤^. *J*(s) is a *p* × *p* matrix, and the rank of *J*(s) is determined by the number of linearly independent neuronal receptive fields in the population (Supplement 4.3). To predict perceptual discriminability from matrix *J*(s), we can eigen-decompose *J*(s). Notice that *J*(s) is a symmetric, positive semi-definite matrix, and all of its eigenvalues are non-negative. Once an image distortion **∊** is along an eigenvector direction of *J*(s), *J*(s)***∊*** = λ***∊***, and the predicted perceptual discriminability is the square-root of the corresponding eigenvalue λ. This is because 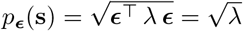.

At a single reference image, two different neuronal response models may predict different receptive field patterns, which results in two different discrimination matrices *J*_1_(s) and *J*_2_(s). To compare between the model-predicted perceptual discrimination, we can compare how much the eigenvalues and eigenvectors of *J*_1_(s) and *J*_2_(s) differ (Berardino et al. [2018]). In the rest of this section, we illustrate such a comparison using a simplified retinal example.

#### Example 1: perceptual distortions with linear retinal cells

In this example, we inspect how perceptual discriminability relates to the receptive field of a population of linear neurons. For simplicity, we consider three distinct cases with populations of two neurons, i.e. one ON cell and one OFF cell.

In the first case (Figure 2A), the ON and OFF cells have the same filter shape, but with opposite signs. Because we assume linear computations, the cells’ receptive fields are identical to their filters. The two cells have linearly dependent receptive fields, so the perceptual discrimination matrix has a rank 1. The most discriminable direction in this case is identical to the shared receptive field direction between the two neurons. In the second case (Figure 2B), the two cells share the same filter location, but the OFF cell has a smaller size. Because the two cells’ receptive fields are linearly independent, the discrimination matrix has rank 2. The second most discriminable direction/perturbation is orthogonal to the most discriminable direction. The second most discriminable direction predicts an additional perceptual sensitivity to contrast, compared to the first population. In the third case (Figure 2C), the filters of the neurons have the same shape but different center locations. The corresponding discrimination matrix is also of rank 2.

**Figure 2:**
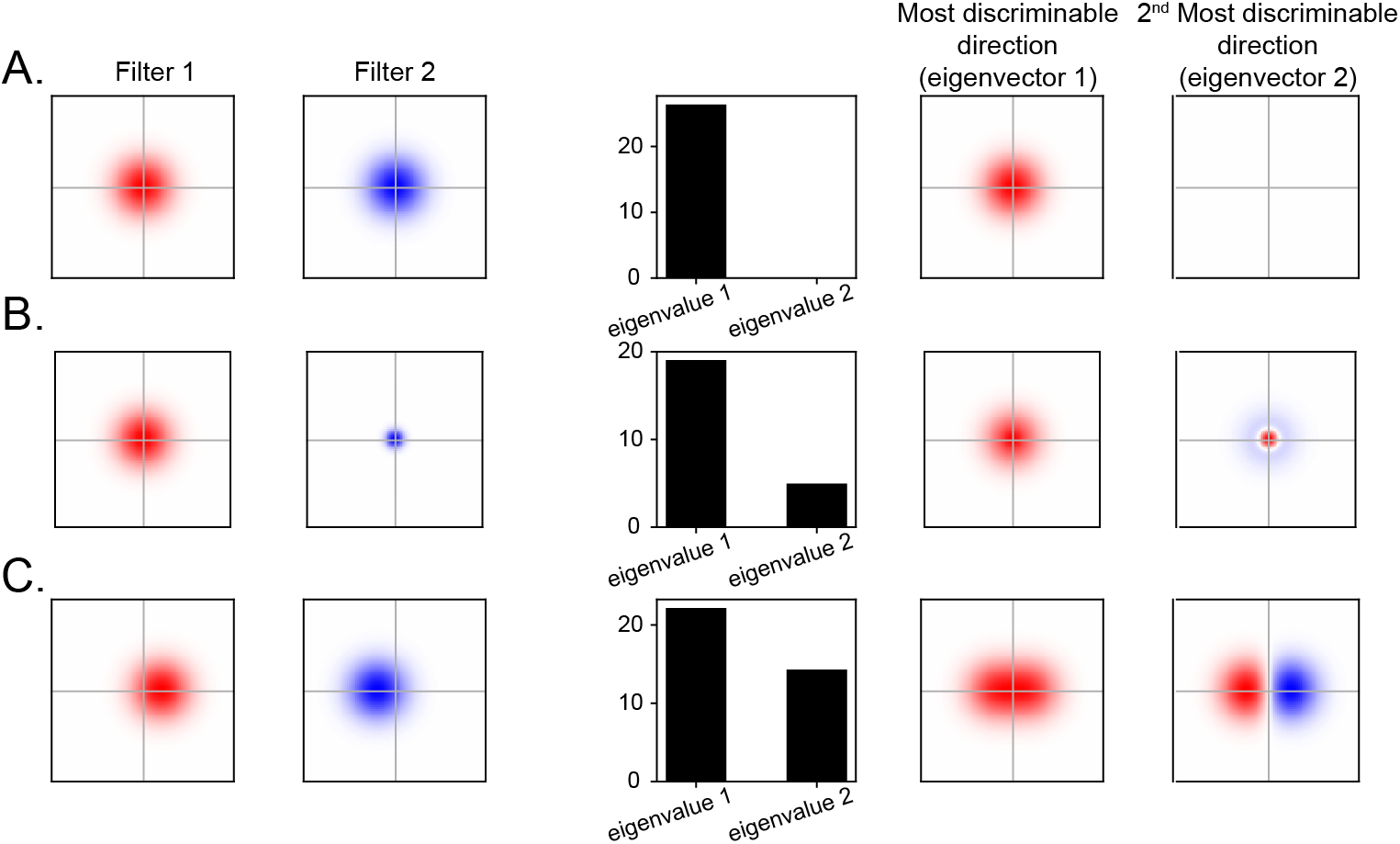
Relating a population’s receptive field to perceptual discriminability. The neuronal population consists of 2 cells, an ON cell and an OFF cell, and both cell types’ image filters consist of a single blob Gaussian (with different signs). *A*. Both the ON and the OFF cells have identical filter but with the opposite signs. Because we assume linear computations for both cells, their receptive fields coincide with their image filters. The two cells have linearly dependent receptive fields, so their predicted effective discriminability has dimension 1, which is reflected by the single non-zero eigenvalue of the discrimination matrix. *B*. The ON and the OFF cell share the same filter location but different filter sizes. Because the two cells have linearly independent receptive fields, they predict a two-dimensional space of effective image distortions. *C*. The ON and the OFF cells do not share the same receptive field location, and they predict a two-dimensional space of effective image distortions.

To summarize, we show in this example that the dimensionality of perceptual discrimination can be different for populations with the same number of neurons. Perhaps more interestingly, even if the neurons have simple and the same overall filter shapes, as a population they can perceptually discriminate complex perturbations.

### 2.2 Static nonlinear neuronal responses

Linear models are not realistic models to describe neuronal responses in general, nevertheless it is a useful start to capture small response changes. This is because typically, neuronal responses smoothly vary with image inputs, and slight image distortions cause small response changes that can be well-approximated by a linear model (Taylor expansion). Empirically, physiologists observed that neuronal responses over a large stimulus range deviate from the linear predictions in at least two ways. First, neuronal responses are often accounted by the number of spikes they fire, and spike count is a non-negative number. Second, neuronal responses saturate. For example, doubling stimulus intensity increases visual neurons’ responses, but typically the extent of increase is less than 100% (e.g. Shapley [1984]). In this section, we expand upon the linear model, by applying a nonlinear transform to the output of the linear computation. Because of this nonlinearity, the intensity of a single neuron’s receptive field may vary across reference images, whereas a linear neuron’s receptive field stays the same for all references. We begin this section by analyzing how this nonlinearity contributes to perceptual discrimination using a single neuron, before we move on to population analyses.

#### How does perceptual discriminability relate to a single linear-nonlinear (LN) neuron’s receptive field?

Suppose a neuron’s response *r* can be summarized using a linear computation followed by a nonlinear function *g*(*x*), where *x* indicates the linear output f^⊤^s:

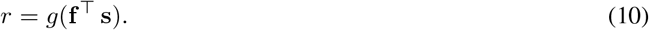

Different forms of *g*(*x*) capture different empirical observations that are inconsistent with the linear predictions. For example, *g*(*x*) can be a soft rectified linear function (e.g. *g*(*x*) = *e^x^*) that eliminates negative linear outputs; or *g*(*x*) can be a sigmoidal function that captures a neuron’s increased response sensitivity at low, and decreased sensitivity at high stimulus intensities (e.g. Brinkman et al. [2008]). The intensity of an LN neuron’s receptive field may vary when assessed using different reference images. To observe this, we first compute the directional derivative of response *r* with respect to reference s along distortion ***∊*** (using the Chain rule):

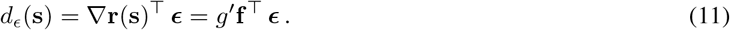

Here, *g*′ is a shorthand for 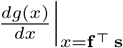, and it is a scalar quantity that indicates the derivative of *g*(*x*) at the linear output *x* = f^⊤^s. An LN neuron’s receptive field consists of the neuron’s image filter f scaled by *g*′: ∇*r*(s) = *g*′f. When reference image s changes, the neuron’s linear output changes, so is the value of the scalar *g*′. The intensity of the LN neuron’s receptive field is determined by *g*′, which generally varies across reference images. But the shape of the receptive field, which is determined by f, stays the same (see Figure 3A). *g*′ is often interpreted as the neuron’s response gain at reference s (e.g. Shapley [1984]). Notice that the response gain depends on the reference, but not on image perturbations ***∊***.

**Figure 3:**
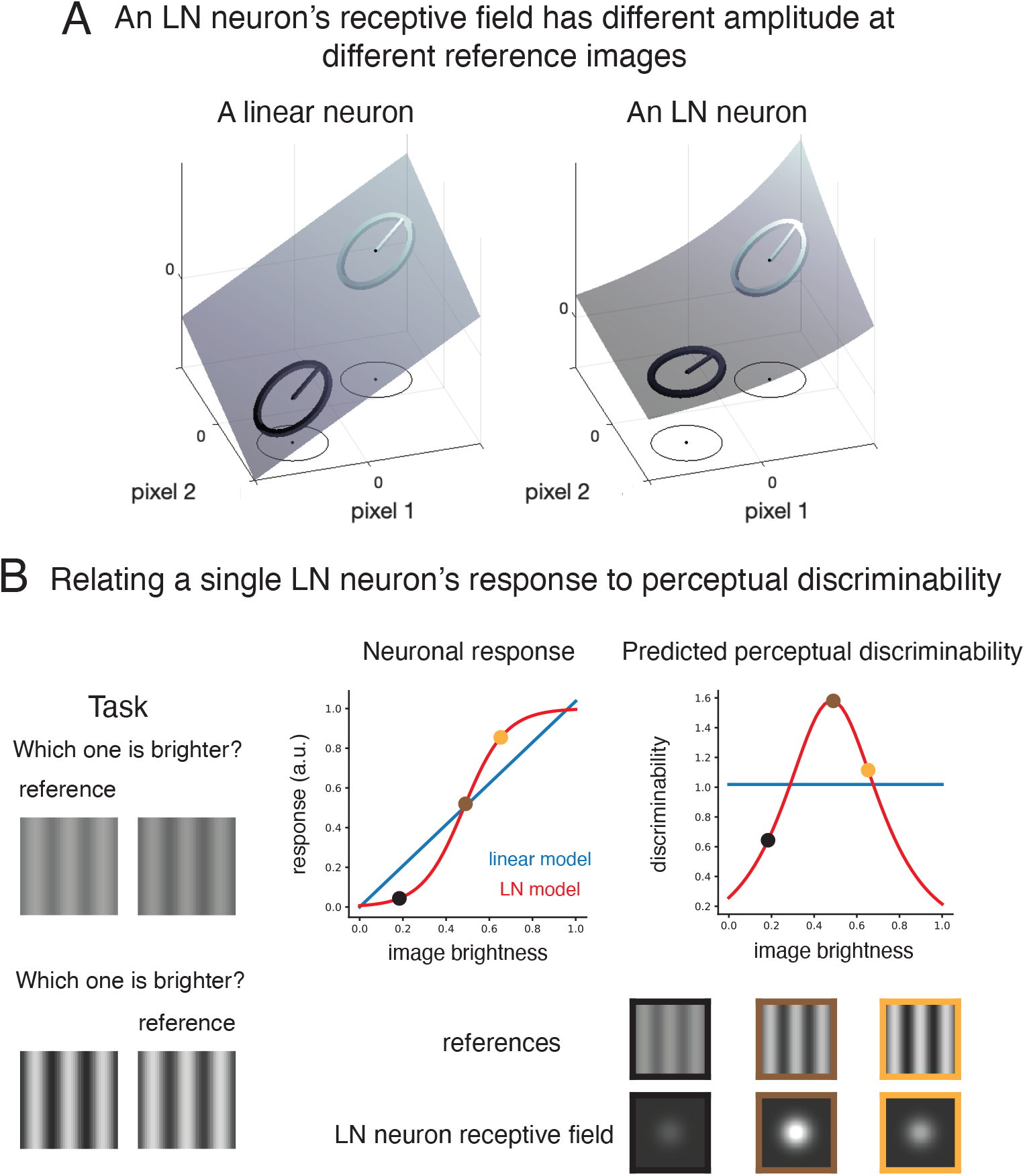
A single neuron’s LN computation and its perceptual consequences. *A*. A linear neuron’s receptive field does not depend on the reference. We plotted two reference images, and equi-distance distortions around each reference. The vector within each circle indicates the linear neuron’s receptive field, and for both reference images, the receptive field vectors share the same length and direction. An LN neuron, on the other hand has the same receptive field pattern across different reference images, and this is reflected by the receptive field vectors at different references share the same direction. However, the length of the receptive field vector may differ at different references. *B*. Suppose both a linear and an LN neuron share an Gaussian image filter. Both neurons were presented with gratings of different intensities. Both neurons were given a task to discriminate between a reference grating, and a grating with an intensity slightly higher than the reference, and this task is repeated for reference gratings of different intensities. The linear and the LN neuron have different response profiles, therefore predict different patterns of perceptual discrimination. The linear neuron has the same receptive field at all references, so its predicted discriminability is the same for all references. The LN neuron’s receptive field, as well as the predicted discriminability differs at different references.

Perceptual discriminability for an LN neuron can be expressed as:

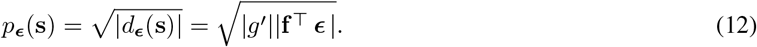

The magnitude of the angular comparison |f^⊤^***∊***|, and the magnitude of the response gain |*g*′| co-determine perceptual discriminability. In general, an LN neuron’s response gain can be positive or negative (±*g*′), but only the magnitude contributes to perceptual discrimination. In the rest of this section, to better appreciate the perceptual advantage of the LN computation compared to the linear computation, we analyzed an example to illustrate how a linear and an LN neuron give rise to different perceptual predictions.

##### Example 2

In this example, we illustrate that a linear and an LN neuron make different perceptual predictions (Figure 3B). First, we assume that both the linear and the LN neuron share a single Gaussian image filter (like the ON cell in Example 1). A set of perceptual tasks were delivered to both neurons. In each task, a grating image (the reference), and a slightly distorted version of the grating were shown to both neurons. The image distortion is along the direction of light increment for all pixels in the reference image. The neurons’ task is to distinguish between the two images, and decide which one is the reference, and which one is the distortion. We repeated the same task for different reference images, and the overall pixel intensities are different for different references.

The two neurons have different response profiles, therefore are making different perceptual predictions. The linear neuron has the same receptive field for all references, and because perceptual discriminability is completely determined by the receptive field, the linear neuron predicts the same level of discriminability at all references. The LN neuron, on the other hand, has different receptive field intensities at different reference images. When the receptive field has high intensity, the neuron predicts a large perceptual change for the image distortion, and when the receptive field is low in intensity, the prediction of the perceived image distortion is small. The nonlinearity in the LN neuron is assumed a sigmoidal form, which has an expression: *g*(*x*) = 1/[1 + *e*^−10*x*+5^].

Overall, the LN neuron’s response gain changes with stimulus intensity, and its perceptual discriminability scales with the (non-negative) response gain.

#### How does perceptual discriminability relate to a population of LN neurons’ receptive field?

Suppose a population consists of *n* neurons, and each neuron has a filter f_*k*_ and a static nonlinear transform *g_k_*(*x*). Both the filters and the nonlinearities are generally different across neurons (e.g. Chichilnisky and Kalmar [2002]), and this LN population’s response can be summarized as:

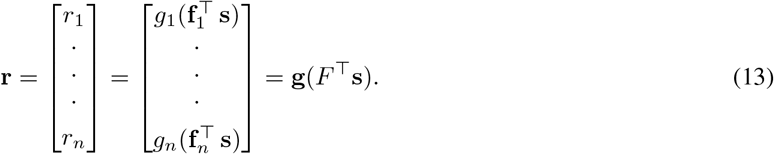

To compute directional derivative for this neuronal population, we first define 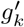 as the derivative of the *k^th^* neuron’s point-wise nonlinear transform with respect to the linear output, 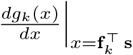. To express the directional derivative as a matrix computation, we further define Γ^s^ as an *n* × *n* diagonal matrix that has 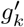 as its *k^th^* diagonal entry:

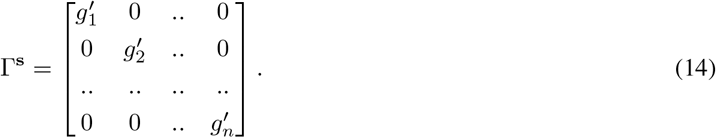

The superscript s of Γ^s^ is to emphasize that the diagonal entries of this matrix can take different values at different reference images. Using this gain matrix Γ^s^, the directional derivative for the LN population can be expressed as:

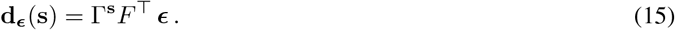

Receptive field for this LN population has an expression ∇*r*(s) = FΓ^s⊤^. Notice that different from a single LN neuron’s perceptual predictions, an LN population’s receptive field generally exhibits different patterns across reference images. This is because the entries of the gain matrix Γ^s^ vary across reference images, and the population receptive field consists of neuronal filters being linearly combined via these varying gains. This difference in receptive fields causes the predicted patterns of perceptual discriminability to also depend on the reference image. To predict perceptual discriminability from an LN population’s receptive field, we compute the norm of the directional derivative:

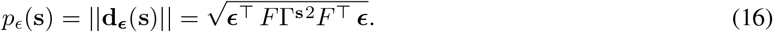

As in linear population analysis, the discrimination matrix *J*(s) = *F*Γ^s2^*F*^⊤^ completely determines the pattern of perceptual discriminability, i.e. the eigenvalues and eigenvectors of *J*(s). Discrimination matrix *J*(s) varies across reference images, because Γ^s^ is dependent on reference image.

At a single reference image, when two neuronal populations have the same receptive field, they predict the same pattern of discriminability. On the other hand, when two populations predict the same discriminability pattern at a reference image, their receptive fields can be different. This is because for any receptive field ∇*r*(s), an orthogonal matrix (i.e. rotation) *Q* multiplied by the receptive field ∇*r*(s) predicts the same pattern of discriminability as ∇*r*(s): ∇*r*(s)^⊤^∇*r*(s) = ∇*r*(s)^⊤^Q^⊤^Q∇*r*(s).

Previously, we analyzed the receptive field of a linear population. Receptive fields of an LN and a linear population are related. For an arbitrary linear population, we can always find an LN population with matching receptive field (Supplement 4.4). In the Supplement, we also show that with a slight relaxation of an assumption, the converse of this statement is true.

In the rest of this section, we use an example to illustrate that nonlinearities play an important role in an LN population’s perceptual predictions. We demonstrate the different patterns of perceptual discriminability predicted by two LN populations that share the same image filters, but have different nonlinear transforms.

##### Example 3

In this example, we analyze two LN populations’ perceptual predictions at a single reference image. The reference image is a single Gaussian blob (Figure 4A). The neuronal population consists of a set of ON filters (truncated Gaussian blobs), and a set of OFF filters (Gaussian blobs identical to the ON filers but with the opposite sign). We endowed the population of ON and OFF cells with two types of nonlinearities, a rectified linear computation (or “ReLU”), or a sigmoidal nonlinearity. Because the reference image consists of light increment only (all pixels are positive), under the linear computation, all OFF cells have responses of value 0 (rectified responses), and some ON cells have relatively high and positive responses (Figure 4B, C, D).

**Figure 4:**
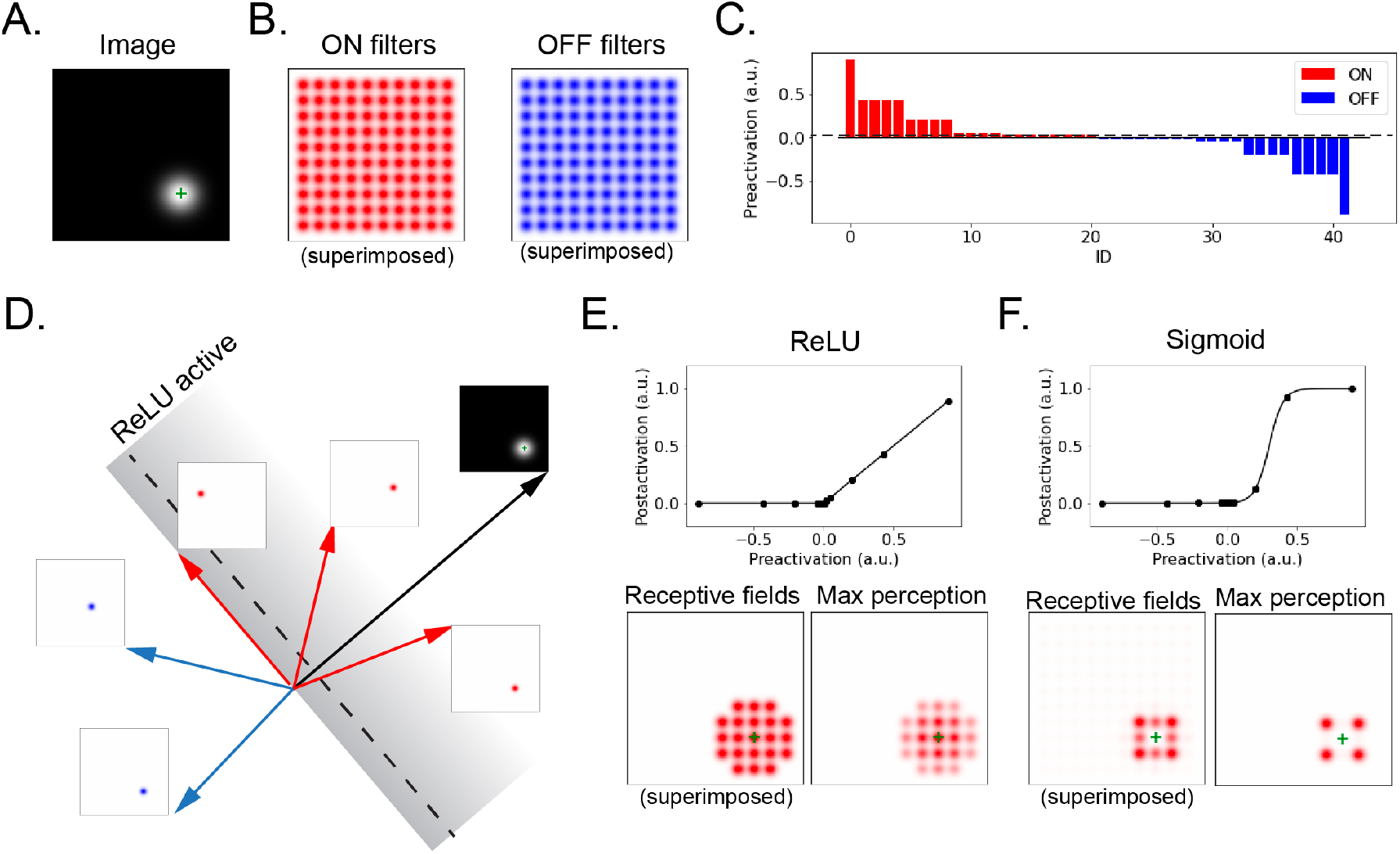
Relating perceptual discrimination to a population of LN neurons. *A, B*. Reference image and a superimposed, i.e. summed, image of the neuronal filters. We used a single Gaussian blob (homogeneous in size) as each ON cell’s filter, and we changed the sign of each Gaussian blob to produce each OFF cell’s filter (blue: positive; red: negative). *C*. We characterize each cell’s linear response (preactivation) profile to the reference image. For this particular reference image, all ON cell’s linear responses are positive, and all OFF cells’ linear responses are negative. The dotted line indicates a threshold for rectification that will be applied to compute the postactivation response. Preactivation values below the threshold will be zeroed in the post-activation and the ones above will be adjusted by subtracting the threshold amount from the preactivation. *D*. A cartoon of rectified linear unit (ReLU) activation. We assumed a thresholded rectification as each cell’s nonlinear transform, and each cell is a ReLU. The dotted line indicates the threshold. *E, F*. We compared the population receptive fields and their implied most discriminable image change under two different nonlinearity assumptions, ReLU and sigmoidal nonlinearities. Compared to ReLU, sigmoidal nonlinearity predicts a different pattern of population receptive field, as well as the most perceivable image change. For all figures here, we assume a threshold that is slightly positive.

The population receptive fields and the patterns of perceptual discriminability are different under the two nonlinear computations. Under the ReLU computation, the predicted population receptive field matches the location of the light increment in the reference image, and the most perceptually discriminable direction also has a similar pattern (Figure 4E). Under the sigmoidal nonlinearity, the predicted receptive fields, as well as the most perceivable direction of image distortion, have patterns that are different from the ReLU predictions ((Figure 4F). This is because the sigmoidal nonlinearity saturates at high level of linear responses, and the neurons have high linear outputs (the image filters that situate at the center of the Gaussian blob in the reference image) tend to have low response gain. The neurons that produce mid-level linear responses tend to have the largest response gain, so that the neurons that situate at the edge of the Gaussian blob in the reference image tend to contribute more to perceptual discriminability.

### 2.3 Cascaded LN computations and neural networks

In previous section, we analyzed the perceptual consequences of linear and LN neuronal computations. LN, compared to the linear model, more closely mimics actual neuronal computations, and its predicted perceptual discrimination starts to resemble our flexible and context-dependent perceptual system. For example, the amplitude of the LN receptive fields, as well as the predicted perceptual discriminability patterns (for a population of LN neurons) vary with reference images. Neuronal computations are generally more complicated than a single layer of LN. How do more realistic neuronal responses, e.g. multiple layers of LN computations, shape our perception? In this section, we examine the receptive fields and perceptual consequences of cascaded LN computations. Empirically, people observed that besides intensity, a single neuron’s receptive field exhibits diverse patterns when measured using different reference images (Goldin et al. [2022]). We illustrate that this empirical observation can be achieved by more than a single layer of LN computation. Overall, more flexible receptive field patterns result in more diverse and richer patterns of perceptual discriminability. In this section, we start with a single-neuron analysis assuming a 2-layer LN computation. Then we move on to a population-level analysis and examine more layers of LNs.

#### How does perceptual discriminability relate to a single 2-layer LN neuron’s receptive field?

First, we consider a single neuron endowed with two layers of LN computations. The neuron of interest responds to the outputs from another set of *m* neurons in the previous layer (called “sub-units”, e.g. Gollisch [2013], Goldin et al. [2022]), and these sub-units take images as inputs (Figure 5A). Each sub-unit has an image filter f_*k*_, and a nonlinear transform *g*_1*k*_(*x*) that follows the linear computation. The subscript 1_*k*_ in *g*_1*k*_(*x*) indicates that function applies to the 1^*st*^ layer *k^th^* neuron. The output neuron (the neuron of interest) sums over m sub-units’ LN outputs, and applies an additional nonlinearity *g*_2_(*x*) to the linear sum. We can summarize this two-layer LN neuron’s computation as:

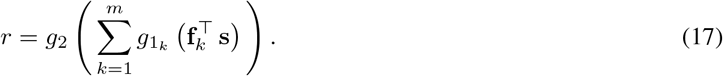

We can compute this neuron’s receptive field to see how its pattern varies with reference images (Figure 5B). We use1 to denote an *m* dimensional column vector with entries 1, and *F* to denote the population filter (*p* × *m*) of the sub-units. The neuron’s receptive field ∇*r*(s) can be expressed as:

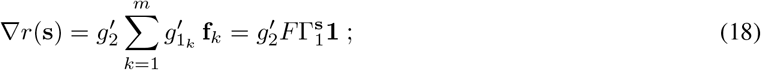

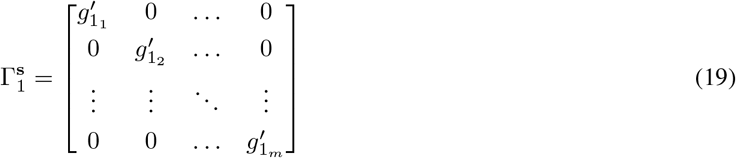

Notice that this neuron’s receptive field is composed of a weighted sum of sub-unit filters {f_*k*_}_*k*_, and the weight for each sub-unit consists of two components, 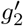 and 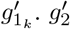 is shared across all sub-units, and modulates how much all sub-units contribute to the output neuron’s receptive field. 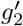 is analogous to the gain modulation in the single-layer LN neuron’s receptive field (Equation 14), The other modulator 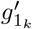 for the two-layer LN neuron is generally different for each sub-unit, and 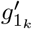 modulates how much the *k*^*th*^ sub-unit filter contributes to the output neuron’s receptive field. The linear output of each sub-unit varies across reference images, and the set of gain modulators {*g*_1*k*_}_*k*_ correspondingly vary, which gives rise to different linear combinations of sub-unit filters in the output neuron’s receptive field. To compute the perceptual discrimination patterns for the two-layer LN model, we can compute the discrimination matrix *J*(s) as before, and *J*(s) =∇*r*(s)^⊤^∇*r*(s). To better appreciate the diversity of receptive field patterns that an 2-layer LN introduces, next we examine an example that compares the receptive field of an 2-layer LN neuron to that of a single-layer LN neuron across different reference images.

**Figure 5:**
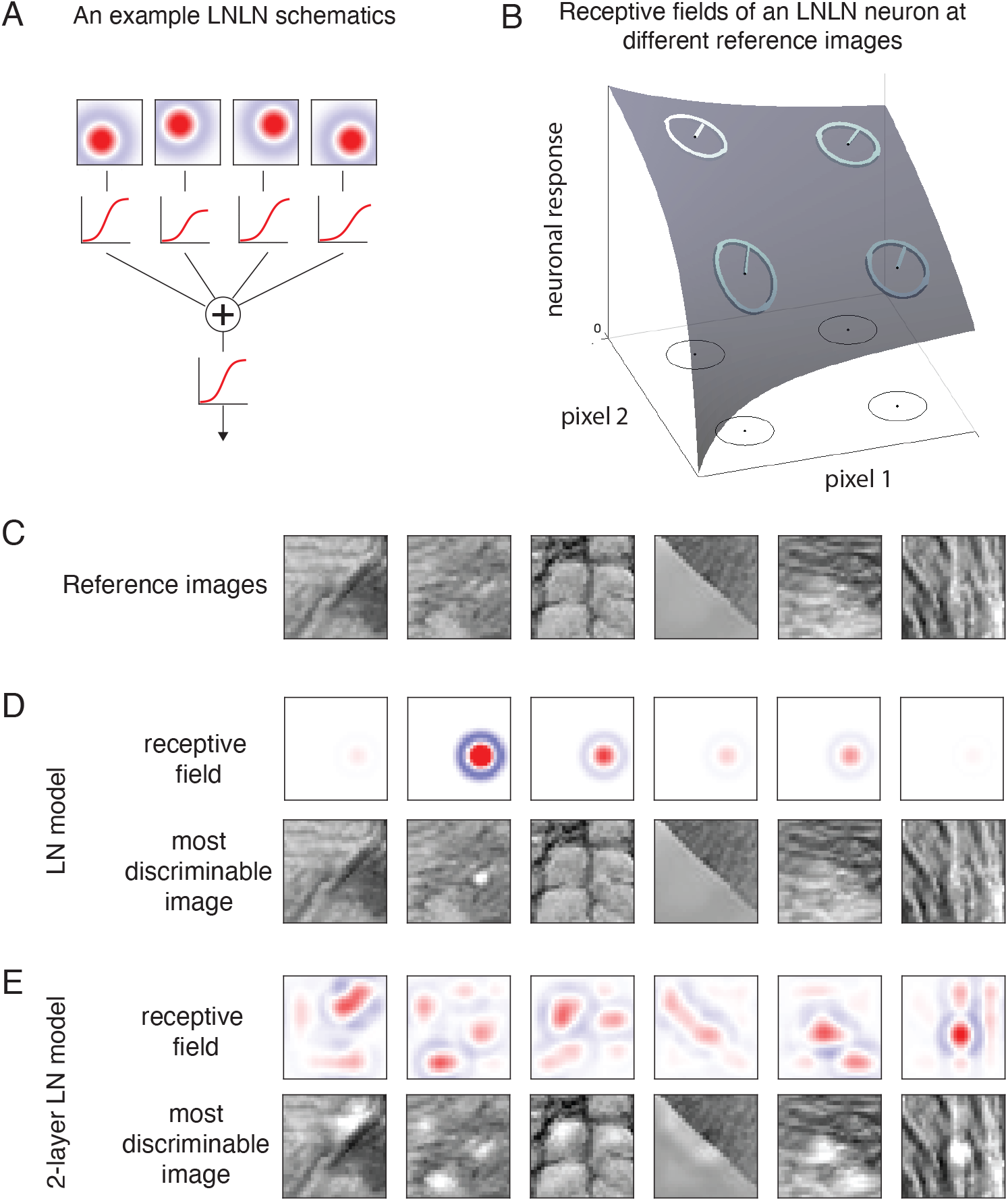
2-layer LN models change receptive field patterns for a single neuron. *A*. Schematics for a 2-layer LN model. B. A cartoon illustrates that under the 2-layer LN model assumption, the receptive field direction (or pattern) may differ for different reference images. C. Using 6 different reference images (natural image patches), we compared a single cell’s receptive field measured under 2 model assumptions, an LN model and a 2-layer LN model. The model implementation details can be found in Supplement, and for all models, the first-layer sub-unit image filters are assumed as a difference between concentric Gaussians with different center locations. D. For the LN model, the receptive fields for different references share the same shape but differ in intensity, the most discriminable image (based on a single neuron’s computation) follows this pattern. E. For the 2-layer LN, the patterns of the receptive field at references generally differ, and reflect some reference image properties (e.g. reflects the regions of high contrast in reference images).

##### Example 4

Here, we examine different receptive field patterns under the assumption of two different neuronal models (Fig. 5C). To illustrate the generality of our conclusions, we sampled six patches of natural images as reference images.

First, we assume an LN computation. Under this assumption, the receptive field of a neuron with center-surround image filter exhibits different intensities at different reference images. The shape of the neuron’s receptive field coincides with the filter and maintains the same across reference images (Fig. 5D). For the 2-layer LN computation, we assume that the output neuron takes inputs from a collection of sub-units with center-surround image filters. The filter shapes of the sub-units lay uniformly across the spatial extent of the image patch. Under this assumption, the receptive field of the output neuron drastically differs both in shape and intensity when measured using different natural images (Fig. 5E). For example, the receptive fields tend to have patterns that correspond to areas of high contrast in the reference image (Goldin et al. [2022]). In both the 2-layer LN and the single-layer LN examples, we illustrate the neuron’s receptive fields, as well as the most perceptually discriminable images, each of which is the neuron’s receptive field added on top of the reference image.

#### How does perceptual discrimination relate to a feed-forward neural networks?

If a neuron or a neuronal population consists of more than two layers of LN computations (e.g. the neurons are deep in the visual cortex), the receptive field patterns of these neurons can be very flexible, and the components of these receptive fields can vary across multiple spatial scales. To make this intuition concrete, we examined cascaded LN model with multiple layers of computations for a neuronal population. The cascaded LN model has the following recursive structure:

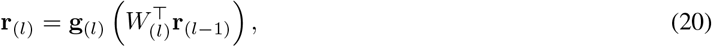

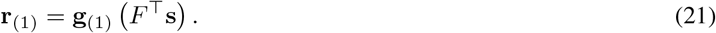

The index (*l*) indicates the layer of the neuronal computation. The 1^*st*^ layer sub-units have filter **F**, and directly take images as inputs. r(_*l*_) represents the *l^th^* layer neuronal outputs, and *g*(_*l*_) is the element-wise nonlinear transform for all neurons in the *l^th^* layer. *W*(_*l*_) indicates the weights used to linearly combine the (*l* – 1)^*th*^ layer neurons’ responses to the *l^th^* layer’s linear outputs. Previously in the 2-layer LN analysis, we assumed a single output neuron, and used a simplified weight matrix *W* = 1, which is a column vector whose elements are value 1, and all sub-unit outputs converge to a single last-layer neuron with equal weight 1.

The Jacobian (transpose of the receptive field) of a population of cascaded LN neurons can be obtained by taking the derivative of the recursive equation:

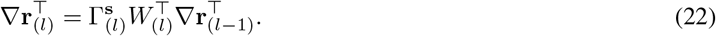

And overall, the receptive field of cascaded LN population is a linear combination of the input-layer sub-unit filters **F**:

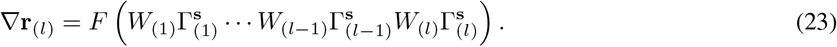

The dimensionality of the receptive field ∇*r*(_*l*_) is limited by the “bottleneck” – the smallest rank of *W*(_*k*_) or 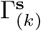 of the network computations, because of the rank property in matrix computations: *rank*(*AB*) ≤ min [*rank*(*A*), *rank*(*B*)]. A small rank of *W*(_*k*_) could be due to the number of neurons in the *k^th^* layer is smaller than that of the other layers. A small rank of the diagonal matrix 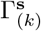 could be due to rectification. For example, most of the *k^th^* layer neurons’ linear outputs are below 0, and the nonlinearity for the *k^th^* layer computation is ReLU with a threshold at 0. In other words, sparse neuronal responses could contribute to the low dimension of perceptually-relevant image distortion, or the low-dimension perceptual representation of images that has been proposed in the literature (e.g. Waraich and Victor [2021],Thompson [1994], O’Toole et al. [1993], Edelman and Intrator [1997]).

### 2.4 Stochastic neuronal responses

In previous sections, we assumed that neuronal responses are deterministic – repeated presentations of a stimulus s always trigger the same pattern of neuronal responses. This is a common assumption made in biological and artificial networks (Duong et al. [2022]). In fact, up to date, most artificial networks are deterministic, albeit achieving incredible capabilities. However, neuronal spiking outputs are stochastic, and perceptual scientists have realized for a long time that the stochasticity in neuronal responses contribute to our stochastic perceptual representation of the world (e.g. Green and Swets [1966], Zhou et al. [2021]). In this section, we show that by making a slight adjustment to our previous analysis, we can flexibly include different types of noise into predicting perceptual discriminations from neuronal responses.

### 2.5 How to relate neurons’ stochastic responses to perceptual discriminability?

In each trial of a perceptual discrimination experiment, an observer is either presented with a reference image s, or its slightly perturbed version (s + *∊*). The observer needs to judge whether they were shown the reference (in which case they answer “No” distortion), or the distortion (in which case they answer “Yes”). When the image distortion is small, often times observers’ answers to the presentation of the same image become stochastic. Classical psychophysics and signal detection theory attribute observers’ stochastic answers to their stochastic perceptual representation: to an observer, what is seen as an distortion in one trial could be seen as the reference in a different trial. It is believed that observers’ stochastic perception is the consequence of stochastic neuronal responses Green and Swets [1966], Kingdom and Prins [2009]. Poisson is a classic model of stochastic neuronal responses: when a single image is presented repeatedly, a neuron can fire different number of spikes, and the spiking variance grows in proportion to the mean.

To quantify the effect of stochastic neuronal responses on perceptual discriminability, we first introduce some new notations. For a population of n neurons, we use *p*(*γ*|s) to denote the response distribution for stimulus s, and *γ* is an n-dimensional response vector. For example, when neuronal responses are distributed as independent Poissons, *p*(*γ*|s) = *Poisson*[r(s)], and each neuron (indexed *k*) may have a different mean response r_*k*_(s) that varies as a function of stimulus. For example, r_*k*_(s) can follow an LN model as we previously analyzed.

In previous sections, we’ve seen that matrix *J*(s) = ∇*r*(s)∇*r*(s)^⊤^ of a deterministic model completely characterizes the pattern of perceptual discrimination. To relate stochastic neuronal responses to perception, we use Fisher information, which has an analogous expression to *J*(s). For deterministic neuronal responses, we obtained the discrimination matrix by quantify the magnitude of change in the stimulus-dependent neuronal responses. Fisher information relates to quantifying the magnitude of change in response distribution (Zhou et al. [2021]), and can be defined as:

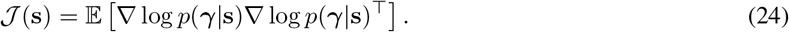

∇ log *p*(*γ*|s)⊤ is the Jacobian of the log of the population response distribution function *p*(*γ*|s) with respect to s, and E stands for taking an expectation over all *γ*. To compute the magnitude of population response distribution change with respect to an image distortion direction ***∊***, we have the following expression analogous to the deterministic analysis (Berardino et al. [2018]):

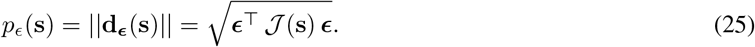

By replacing *J*(s) for deterministic neuronal responses by Fisher information 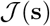, we incorporated neuronal response noise to account for perceptual discrimination. For more details and intuitions of Fisher information, see Seung and Sompolinsky [1993], Itti et al. [1997], Averbeck and Lee [2006], Paradiso [1988], Series et al. [2009], Solomon [2007], Moreno-Bote et al. [2014], Kanitscheider et al. [2015]. For arbitrary distributions, Fisher information is generally difficult to compute, and in practice, a lower bound of Fisher information (Stein et al. [2014]), which only involves the mean (r(s)) and the covariance (Σ(s)) of the response distribution, has often been used (e.g. Beck et al. [2011]):

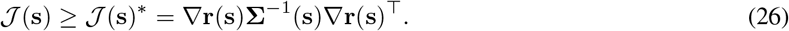

For many common distributions (including independent Poisson distribution), this lower bound is exact (Zhou et al. [2021]). Notice that the lower bound expression 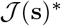 is in a quadratic form, similar to the expression of *J*(s) (*J*(s) = ∇*r*(s)∇*r*(s)^⊤^), with an additional Σ^−1^(s) that transforms the spike rate representation of an image distortion. When we perturb a reference image along an equi-distance circle in the image space, first, the mean neuronal responses (the spike-rate image representation) compresses the image perturbations along some directions, and expands them along some other directions, forming an elliptical representation of the image distortion (Figure 6A). Moreover, this spike-rate elliptical representation is further transformed by the inverse of the covariance matrix of neuronal response, and this final elliptical distortion is the stochastic representation of the distortions in the image space. To gain an intuitive understanding of how stochastic and deterministic representations differ, in the rest of the section, we examine perceptual discriminability predicted by both models using a common set of reference images.

**Figure 6:**
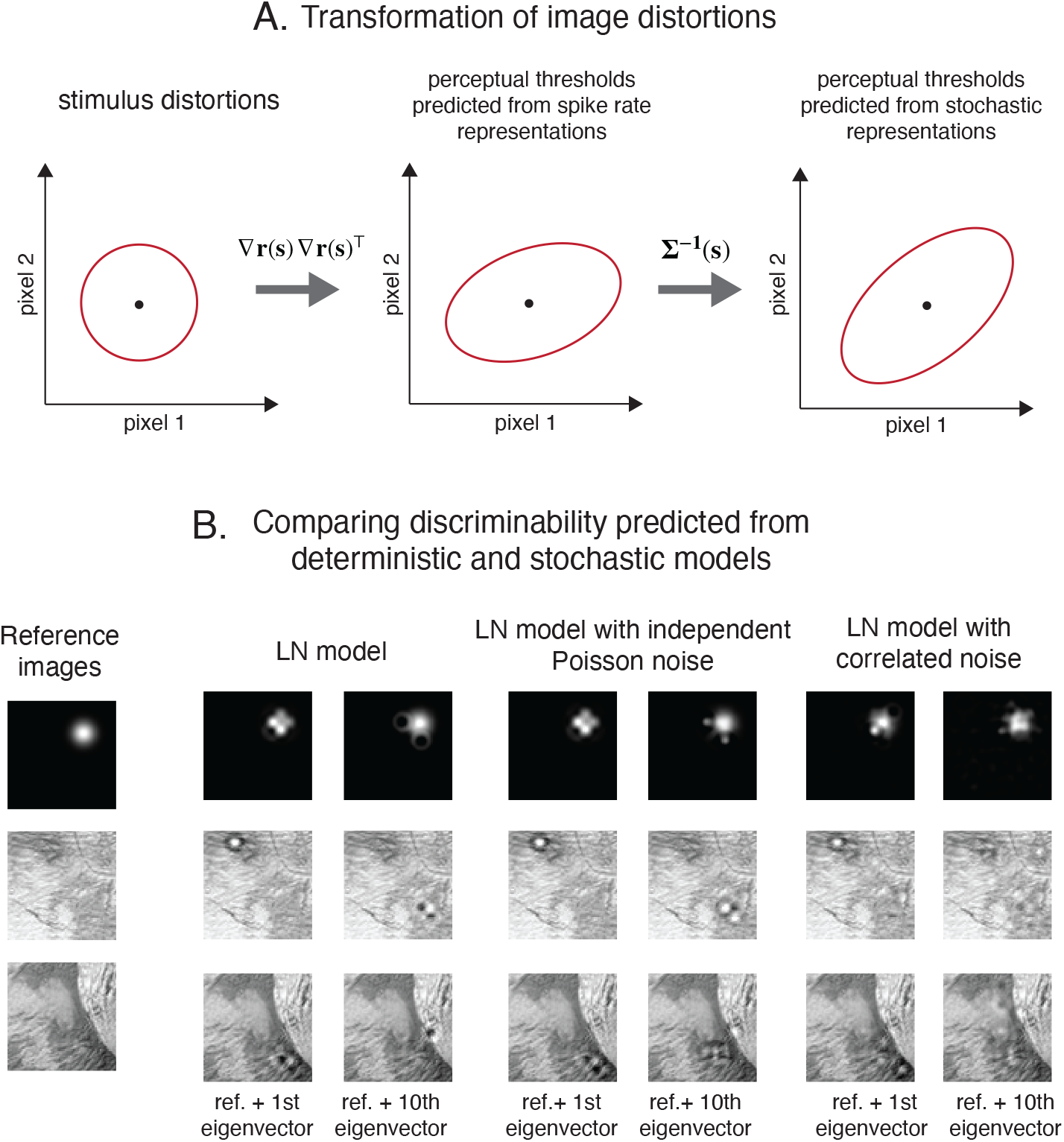
Population receptive fields and perceptual discrimination predicted using deterministic and stochastic models. *A*. The two-stage transformations from stimulus perturbations to perceptual thresholds predicted by stochastic representations. The perceptual thresholds are the discriminabilities projected back to the image space, and shares the same unit as the image. For how the geometry of perceptul thresholds relate to that of discriminability, see Supplement 4.5. *B*. Comparing between deterministic and stochastic LN models’ perceptual predictions. We used 3 references images, an artificial reference image (a single Gaussian blob), together with two natural image patches. For all following panels’ model computations, we assumed a population of concentric difference-of-Gaussian ON-center image filters, and a population of OFF-center image filters. First we assumed an LN model, and compared two images: one perturbed in the first eigenvector direction (corresponding to the largest eigenvalue), and other one perturbed in the 10^*th*^ eigenvector direction. We performed the same analysis for the LN model assuming independent Poisson noise, and for the LN model assuming correlated neuronal noise. The predicted perceptual discriminabilities from the third model (correlated neuronal noise) are more distinct compared to the first two models.

#### Example 5

In this example, we use three reference images to demonstrate the perceptual predictions of three different models (Figure 6B). The first reference image is a Gaussian blob (an artificial image), and the other two are naturalistic image patches. Using these three references, we compare across 3 models, an LN model, an LN model assuming independent Poisson noise, and an LN model assuming correlated neuronal noise.

For each model, we assumed a collection of center-surround ON- and OFF-image filters, and examined the perceptual predictions. To compare between models, we computed each model’s predicted image distortions along the first eigenvector direction (the most discriminable direction), and the 10th eigenvector direction (a supposedly lesser discriminable direction). This is not a complete comparison, and here, we select two directions only for the purpose of illustration.

The LN model well-captures perceptual discrimination to the artificial image: the most discriminable image distortion is much more perceptually prominent, compared to the image distortion along the 10th eigenvector. For the naturalistic image patches, however, the LN model’s predicted perceptual discriminability for the first eigen-vector direction and the 10th eigen-vector direction do not match our perceptual experience. This is because both distortions are similarly prominent to our eyes.

The LN model assuming independent Poisson noise makes similar perceptual predictions to the deterministic LN model. In general, neuronal response variance alone may not contribute much to us perceiving changing image patterns, but it can play an important role in perceiving intensity changes (Zhou et al. [2021]). The LN model assuming correlated neuronal noise predict more distinct patterns of perceptual discriminability, compared to the first two models. For this particular pattern of noise correlation that we chose, this model’s predicted most discriminable image distortion seems to be more perceptually prominent than the 10^*th*^ most discriminable image distortion.

## 3 Discussion

In Visual Neuroscience, the definition of receptive field has gone through changes over time. Initially, when people found that visual neurons in retina respond to light increment at certain spatial locations, receptive field referred to the spatial location that a neuron prefers (e.g. Hartline [1938], Kuffler [1953]). Later on, when people learned that even at the same spatial location, visual neurons in the primary visual cortex are sensitive to some image patterns more than others (e.g. Hubel and Wiesel [1959, 1962]), receptive field referred to a neuron’s most preferred image pattern. This definition lasts until these days, and it is the most common description of receptive field in the experimental literature. However, this global definition of receptive field has the limitation that once we go beyond the visual front end (e.g. retina, LGN and V1), finding the global receptive field of a neuron becomes very challenging, and possibly not very useful for current perceptual studies. A neuron deep in the visual cortex may have very complex response patterns (which is generally highly non-convex), and finding the image pattern that triggers the global maximal response is computationally and experimentally challenging. Moreover, it is hard to perceptually interpret an image pattern that globally triggers the maximal response in a neuron. This is because at this point in time, we only have theoretical and statistical tools to understand local perceptual perturbations, and the relationship between global perceptual perturbation and neuronal response is under-investigated.

In this paper, we made a clear distinction between a neuron’s image filter and its receptive field. We used a neuron’s image filter as a computational assumption – the set of parameters to be estimated for the neuron’s first-stage linear computation. We used a neuron’s receptive field to refer to the gradient of its response (or Jacobian for population responses) at a reference image – the perturbation to the reference image that maximally increase the neuronal response. A neuron can assume complicated computations, and sometimes does not have a single well-defined image filter (as in the case of the 2-layer LN model), but we can always measure the neuron’s receptive field (e.g. use spike-triggered average) as long as the neuron’s responses smoothly vary with image inputs.

In our analyses to relate perceptual discriminability to neuronal response change, we assume discriminability is the *L*^2^ norm of the vector that describes the response change. *L*^2^ norm is a natural choice, because it quantifies the Euclidean distance between the neuronal responses to a reference, and to a distorted image. At a reference image, we locally approximated the neuronal response change using a linear model (the directional derivative), and the Euclidean distance is a natural quantification of the distance traversed on this linear hyper-plane, because a smooth response manifold is locally Euclidean. Besides that it is natural, the *L*^2^ norm is also supported by empirical evidence. Poirson and Wandell [1990] and Knoblauch and Maloney [1995] drew a circular perturbation of a reference image in a multi-dimensional stimulus space, and examined whether the corresponding perceptual thresholds follow an elliptical pattern. When perceptual thresholds along the stimulus perturbation directions form an ellipse (or ellipsoid), it is in support of the *L*^2^ norm. If the perceptual thresholds are not elliptical, it would be an evidence in support of perception transforms neuronal representations via other *L^p^* norm (Figure 7). Both papers failed to reject the *L*^2^ norm hypothesis.

**Figure 7:**
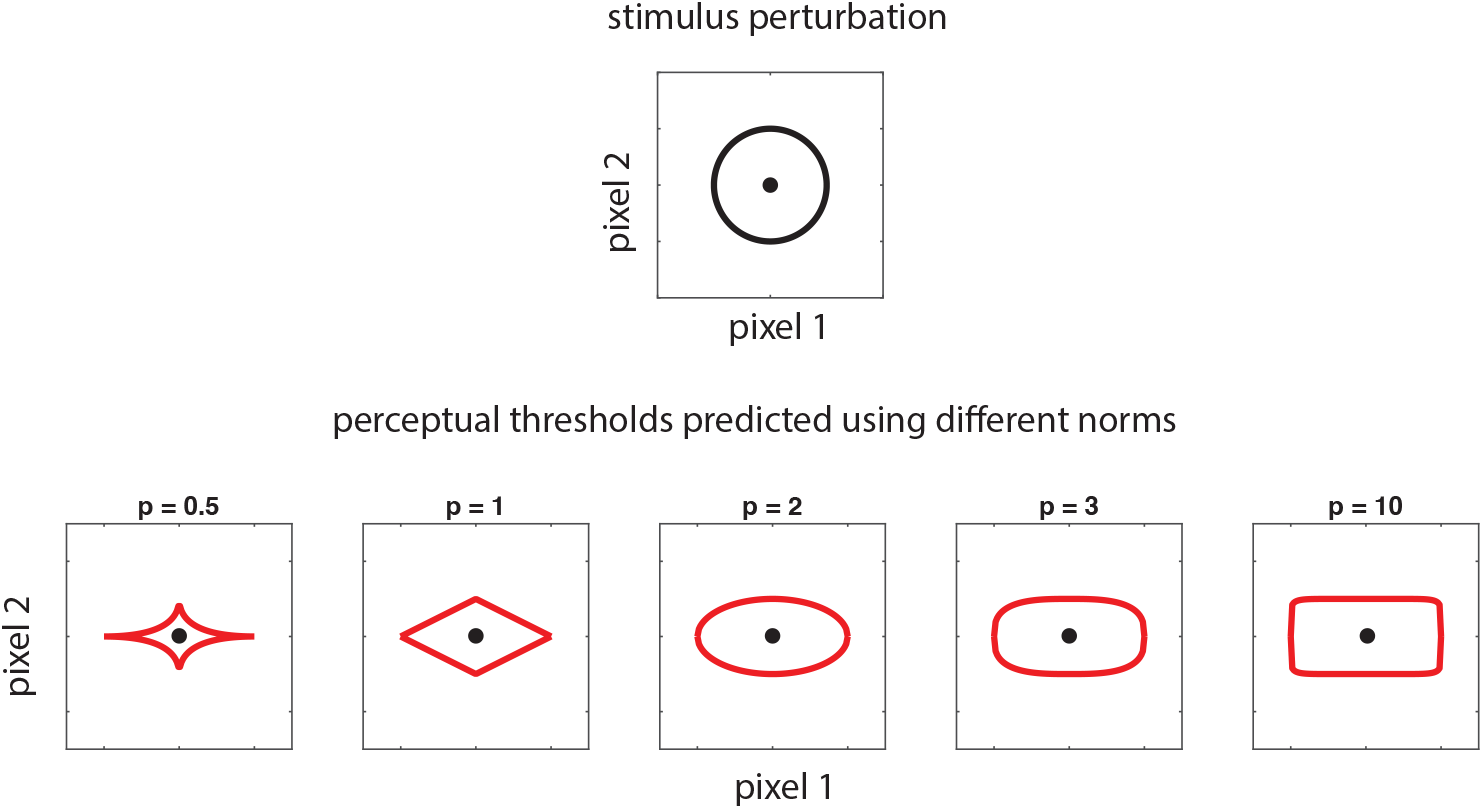
*L*^2^ versus other *L^p^* norms in transforming neuronal responses to perception. In the main text, we assumed perceptual discriminability relates to neuronal response change via an *L*^2^ norm. This is an assumption that sustained empirical tests (Poirson andWandell [1990], Knoblauch and Maloney [1995]). For a circular distortion in the image space, if perceptual change is the *L*^2^ norm of some neuronal computations, the measured perceptual threshold (projected back in the stimulus space) is going to be elliptical in shape. If perceptual discriminability relates to neuronal response change via other *L^p^* norms, the measured threshold along the circular image distortion would form other shapes. For geometrical connection between perceptual thresholds and discriminability, see Supplement 4.5.

Connecting neuronal responses to perceptual discrimination is not a new idea, and there has been at least two different literature that attempt to address this connection. The first one was a theoretical attempt, and has only been recently connected to experimental measures. In this literature, the researchers calculated Fisher information (as a prediction for perceptual discriminability) from a population of independent, or correlated neurons (e.g. Paradiso [1988], Seung and Sompolinsky [1993], Averbeck and Lee [2006], Kanitscheider et al. [2015]), and Fisher information was typically computed along a single stimulus dimension, and sometimes at a number of different reference images. The second literature has an empirical origin (e.g. Spinelli [1966], Britten et al. [1992], Zohary et al. [1990], Liu and Newsome [2005]). This literature made observations over different stimulus domains that discriminability computed for individual neurons may span a large range, and some neurons’ discriminability are comparable to, or even exceed the animal’s perceptual discriminability. In this literature, both perceptual and neuronal discriminability were typically examined using stimulus distorted along a single direction, whereas in this paper, we focus on the analysis using high-dimensional stimuli and distortions. In general, examining perceptual discriminability and neuronal responses using multiple references, and multiple image distortion directions could be data-demanding. Nevertheless, with improved computing power and large scale data-collection effort, this challenge will be alleviated over time. With the surge of studies on (deep) neural networks, the computational tools developed to relate multi-dimensional perceptual distortions to neural responses can bypass the data challenge and still be useful in comparing representations across neuronal models.

## 4 Supplement

### 4.1 Directional derivatives

To understand directional derivatives, first we start with *partial derivatives*, which is the simplest generalization from a single-dimensional derivative calculation. We use e_*j*_ to denote a vector of length *p*, with the *j^th^* entry being 1, and all other entries being 0: e_*j*_ = [0,0,…, 1,…, 0]^⊤^. A partial derivative of the neural response for a given reference image s accounts for the response change when the image is changed along one of its coordinates e_*j*_. Definition of partial derivative takes the following form:

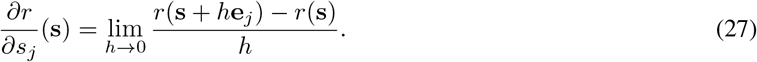

Summarizing partial derivatives along all coordinates {e_*j*_} into a column vector, and we obtain the gradient ∇*r*(s):

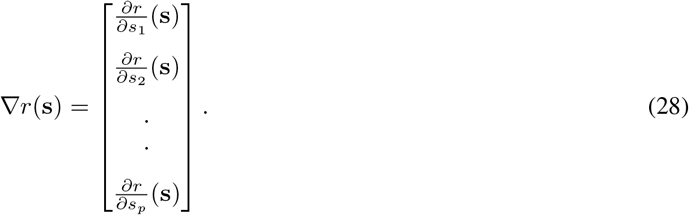

The gradient is an informative summary of local neuronal response properties. First, it provides a linear approximation to the local neuron’s response at stimulus s. When the response function gets smoother, the linear approximation gets better. Any arbitrary point x on this linear hyper-plane (the linear approximation to the response function at a reference image) can be expressed using the gradient:

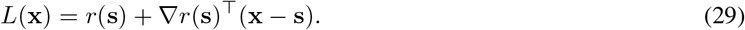

The response change is the largest when an unit length perturbation (x – s) is along the direction of the gradient, so the gradient indicates the direction of image change that triggers the largest response increment.

A *directional derivative* captures the neuronal response change with respect to a change of the image s along an arbitrary direction *∊*. Notice that different from partial derivative, *∊* does not have to be a coordinate direction. Usually, *∊* is also assumed to be a unit vector (‖*∊*‖ = 1) for simplicity. Here is the definition of a directional derivative:

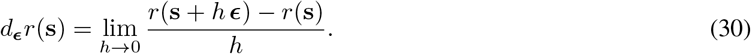

Now we explore the connection between the gradient and the directional derivative. First, we notice that from Eq. 29, response increment is the largest when image distortion *∊* ∝ (x – s) is along the gradient direction. So intuitively, *d*_*∊*_*r*(s) – the extent of response change when s is perturbed along *∊*, relates to the angular comparison between ∇*r*(s) and *∊*. Now we make this intuition concrete. Notice that in Eq. 29, any point x on the linear hyperplane can be expressed using *L*(x), so is (s + *h∊*) for an arbitrary *h*. Now we re-write the directional derivative using function *L*:

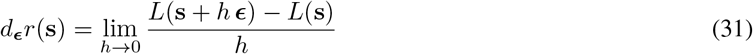

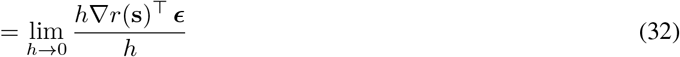

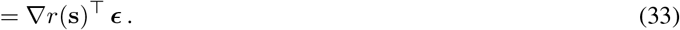

Now we arrived at an expression that connects directional derivative to the gradient.

### 4.2 Supplementary proofs for a linear neuronal population

In this section, we prove two statements in the main text, one is about parallel population filters, and the other is about orthogonal filters.

#### Parallel filters

We prove the following statement: for a population of parallel filters (*F* = [f_1_, f_2_,…, f_*n*_] and f_*i*_, = f_*j*_ for all *i* and *j*), when a reference image s is distorted along an arbitrary direction ***∊***, the population response always adjust along the same direction, but the adjustment can take different amplitudes.

*Proof*. When reference image s is distorted along direction *∊*, change in the population response can be summarized using

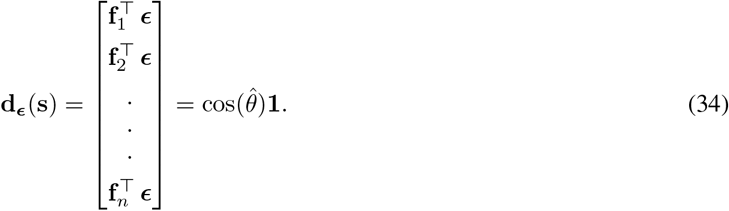

1 is an (*n* × 1) column vector, and 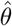 is the angle between filter f_*i*_, and *∊*. For arbitrary distortion direction *∊*, the amplitude of the population response 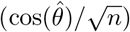 changes, but the population response always adjusts along direction 1.

#### Orthogonal filters

We prove the following statement: for a population of *p* orthogonal filters (and *p* is the number of image pixels), when a reference image s is distorted along arbitrary direction *∊*, the population response adjust along different directions but the adjustment always has the same amplitude.

*Proof*. The population filter *F* in this case is an orthogonal matrix, and the amplitude of the population adjustment has the following expression:

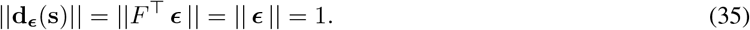

So the population adjustment has the same amplitude for different image distortions.

### 4.3 The rank of the discrimination matrix *J*

Here, we show that the number of linearly independent neuronal receptive fields within a population determines the rank of the discrimination matrix.

First, we demonstrate this by assuming a linear population, the population filters of which can be indicated using *F*. The discrimination matrix *J*(s) can be written as *FF*^⊤^, which can be further expressed as a sum of outer-products of individual neurons’ image filters:

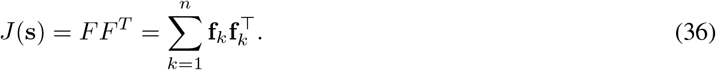

Each item within the summation, 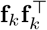 is a rank-1 matrix, and the rank of the overall sum is determined by the number of linearly independent image filters, and is at most *n*, because the population consists of *n* neurons.

Similarly, if we have a nonlinear population, *J*(s) = ∇*r*(s)∇*r*(s)^⊤^, and a similar analysis applies. Instead of summing up the outer-product of individual image filters, we sum up the outer-product sof the gradient of each neuron’s response.

#### Proof for Eqn. (36)

Let e_*k*_ define an *n* dimensional vector whose *k^th^* element is 1 and 0 everywhere else. Then f_*k*_ = *F*e_*k*_. Therefore,

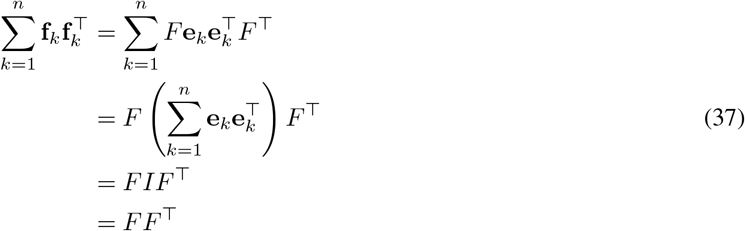

where *I* is the *n* × *n* identity matrix.

### 4.4 How does LN population receptive fields connect to linear population receptive fields?

We show that at a single reference image, we can always find an LN population that has an equivalent receptive field as an arbitrary linear population.

For a linear population with filter *F*, its predicted perceptual discriminability is determined by *J*(s) = *FF*^⊤^ · *J*(s) is a symmetric and positive semi-definite matrix, and can be eigen-decomposed as *J*(s) = *Q*Λ*Q*^⊤^, where *Q* is an orthogonal matrix, and Λ is a diagonal matrix with non-negative eigenvalues. We define *Q* as the LN population’s filters, and Λ^½^ as the gain matrix. Then the receptive field of the LN population, ∇*r*(s) = *Q*Λ^½^, is the same as the linear population’s receptive field *F*.

Now let’s show the converse of this statement. In the main text, we assumed the length of each neuron’s image filter f_*k*_ is 1: ‖f_*k*_‖ = 1. Here, the exact converse statement is false. But we can prove the converse statement under the relaxation that all image filters have length *c*: ‖f_k_‖ = *c*, and *c* is an arbitrary positive number. We show that under the relaxed filter norm assumption, for any arbitrary LN population, we can find a linear population with a matching receptive field.

For any arbitrary LN population, the discrimination matrix *J* can be eigen-decomposed as *Q*Λ*Q*^⊤^. To find the corresponding linear population with matching receptive field, we need to find an orthonormal matrix *R* such that

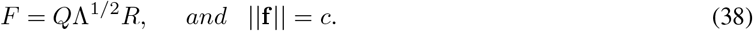

Note that *J* = *FF*^⊤^ since *RR*^⊤^ = *I*.

The norm condition above is equivalent to

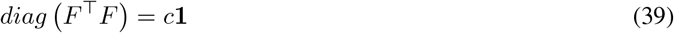

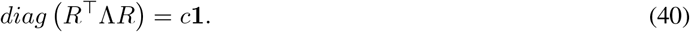

We prove that the above equality can be satisfied for a certain construction of *R* and *c*. Consider the case where the columns of *R*^⊤^ are the harmonics of sine and cosine waves of norm 1:

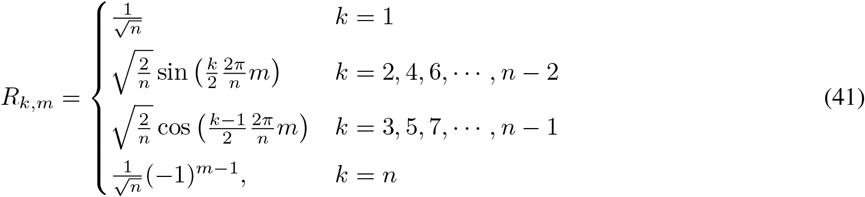

The matrix *R* is visually illustrated in Fig.8. In this case, *R*^⊤^Λ*R* is always a circulant matrix for any arbitrary diagonal matrix Λ (Fig. 8). It is a known property that circulant matrices can be orthogonalized by Fourier basis, and also a symmetric circulant matrix can be created with *R*^⊤^Λ*R* where *R* is a matrix with its rows as sine and cosine waves as defined above (Golub and Van Loan [2013]). Since the diagonal of a circulant matrix is uniform, the norm condition ‖f‖ = *c* is satisfied. In short, with the above *R*, the resulting *F* will have the same norm for its columns, regardless of what *Q* and Λ are.

**Figure 8:**
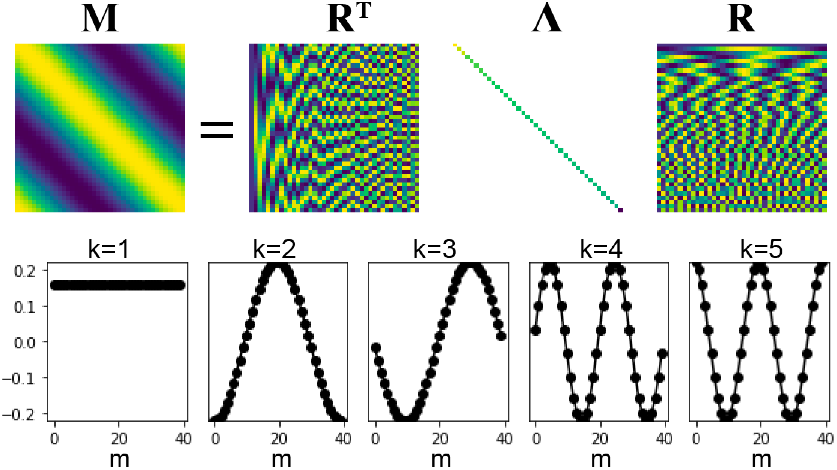
All and only symmetric circulant matrices (e.g. the matrix *M* in the figure) can be decomposed into Fourier basis matrix (R) and a diagonal matrix with non-negative entries (Λ). The bottom plots show the values of the elements in the rows of *R*. The rows are indexed by *k*, and the columns are indexed by *m*.

### 4.5 Geometry between perceptual threshold and discrimination

In this section, we show that if discriminability follows an *L*^2^ norm of change in neuronal responses, the corresponding prediction for perceptual threshold can also be summarized using the *L*^2^ norm.

In the main text, we showed that assuming *L*^2^ norm between neuronal responses and perceptual discriminability, we can use a matrix *J*=∇*r*(s)∇*r*(s)^⊤^ to summarize discriminability. Matrix *J* is symmetric, and positive semi-definite, and geometrically it can be represented using a (high-dimensional) ellipsoid. The longest direction of the ellipsoid is along an eigenvector of J that corresponds to the largest eigenvalue (λ_1_). If image perturbation *∊* is along this direction, the corresponding perceptual discriminability 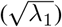 is the largest among all perturbation directions.

Perceptual threshold is the inverse of discriminability. When discriminability is the largest, the perceptual threshold along the same direction (1/λ_1_) should be the smallest among all perturbation directions. To summarize perceptual threshold along all perturbation directions, we are looking for a matrix *J*(s)* such that the eigenvalues of *J*\* is the inverse of the eigenvalue of *J*, 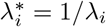. Here, we show that the matrix *J*^−1^ (the inverse or the pseudo inverse of *J*), is the matrix *J*(s)* that we are looking for.

Because *J* is symmetric and positive semi-definite, we can eigen-decompose *J* as *J* = *Q*Λ*Q*^⊤^, such that the columns of *Q* are orthogonal, and Λ is a diagonal matrix with non-zero diagonal entries 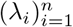. The (pseudo-) inverse of this matrix has a form *J*^−1^ = *Q*Λ^−1^*Q*^⊤^, such that Λ^−1^ has non-zero diagonal entries 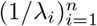.

### 4.6 Glossary of notations

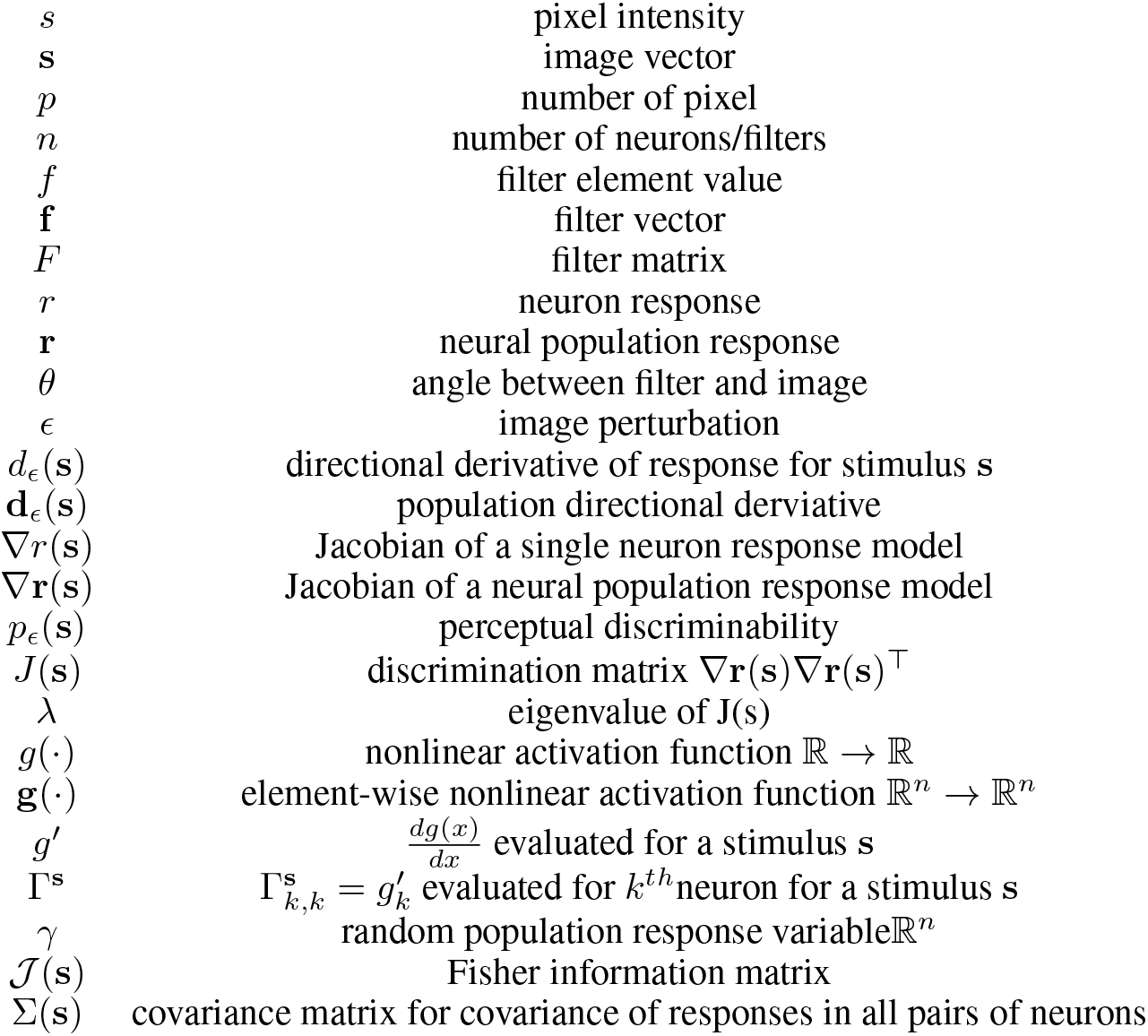

